# CD47 blockade augments anti-GD2 driven phagocytosis *in vitro* but fails to improve *in vivo* efficacy in immune competent, chemoresistant neuroblastoma preclinical models

**DOI:** 10.64898/2026.07.02.736004

**Authors:** Courtney Himsworth, Thomas Jackson, Chantelle Bowers, Sophie Munnings-Tomes, Gaya Nair, Henrike Muller, Elizabeth Tucker, Amy K. Erbe-Gurel, Paul M. Sondel, Louis Chesler, Robbie Mazjner, John Anderson

## Abstract

CD47 delivers a dominant “Don’t Eat Me” signal that inhibits macrophage-mediated clearance of tumour cells. Using immune competent, chemoresistant neuroblastoma (NB) models, we tested a Fc-silent CD47 blocker (ALX301) with anti-GD2 antibody alone and in combination with a clinically aligned temozolomide/irinotecan chemoimmunotherapy backbone. Tumours expressed GD2 and CD47, and bound ALX301. In macrophage coculture assays, anti-GD2 antibody induced dose-dependent phagocytosis, whereas ALX301 or an anti-CD47 antibody alone did not. CD47 blockade in combination with a suboptimal concentration of anti-GD2 antibody showed an additive effect on phagocytosis *in vitro*. *In vivo*, however, ALX301 failed to improve tumour control or survival when added to anti-GD2 or to chemoimmunotherapy in two models. Toxicity was acceptable, showing only mild, expected red-cell changes without organ injury. This form of CD47 inhibition is therefore mechanistically active *in vitro* but insufficient to enhance anti-GD2 antibody-based therapy in immune competent mice bearing a chemoresistant NB, highlighting the potential need for myeloid-reprogramming partners.

## Introduction

Neuroblastoma (NB) is the most common extracranial solid tumour of infancy and early childhood, and outcomes for high-risk disease remain poor despite intensive multimodality therapy. Immunotherapy aims to engage host immunity to recognise and eradicate tumour cells. Anti-GD2 monoclonal antibodies, directed against the widely and highly expressed NB antigen disialoganglioside GD2, are now clinically implemented. When paired with chemotherapy such as temozolomide/irinotecan or cyclophosphamide/topotecan, this chemoimmunotherapy approach improves progression-free survival at relapse versus historical cohorts and has become the platform for rational combinations in ongoing trials(1–6).

CD47 is an ubiquitously expressed transmembrane receptor of the immunoglobulin superfamily(7,8). Its principal ligand, Signal Regulatory Protein alpha (SIRPα), is found on myeloid cells (monocytes, granulocytes, dendritic cells and haematopoietic stem cells) and delivers an inhibitory “self” signal(9). This function was first established on erythrocytes, for which loss of CD47 marks senescent red cells for clearance by red-pulp macrophages(10). Across many human cancers CD47 is overexpressed, prompting development of CD47-SIRPα blocking agents (antibodies and SIRPα-based binders); comprehensive reviews summarise the preclinical and clinical experience(11–14). Given the natural expression of CD47 on haemopoietic cells, Fc-competent anti-CD47 agents may also effect haematological toxicity through off-tumour opsonisation. ALX301 was therefore designed as a Fc-silent CD47 blocker, comprising a high affinity SIRPα D1 domain (engineered for increased binding to mouse CD47) fused to mouse IgG1 Fc with point mutation N297A to abrogate Fcγ receptor engagement while preserving CD47 binding (Fig. 1a).

**Figure 1.**
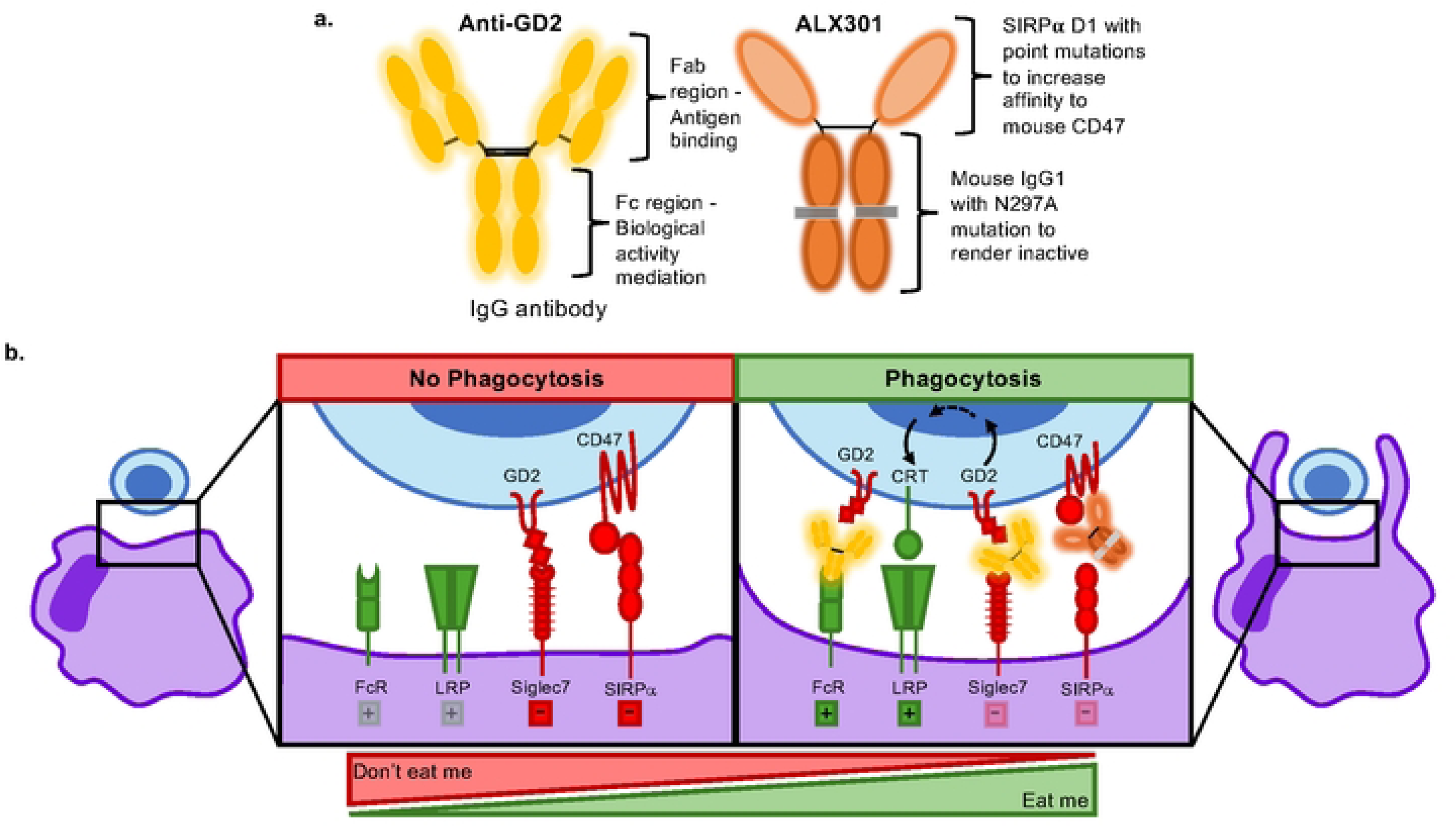
Mechanism of phagocytosis induction by macrophage-targeted immunotherapies. **a.** Schematic of anti-GD2, a conventional monoclonal antibody, and ALX301, a fusion protein molecule. **b.** Overview of signalling involved in phagocytosis regulation. Anti-GD2 and ALX301 treatments shift signalling to favour macrophage-mediated tumour cell phagocytosis.

Macrophages clear antibody-opsonised targets through Fcγ receptor-mediated antibody-dependent cellular phagocytosis (ADCP), if pro-phagocytic “Eat-me” cues, (e.g. calreticulin), outweigh inhibitory “Don’t Eat Me” signals including CD47. Blocking CD47 can tip this balance toward phagocytosis(15). At the tumour-macrophage synapse, anti-GD2 provides the opsonising Fc to engage Fcγ receptors. Anti-GD2 also blocks GD2 binding to its inhibitory ligand, Siglec-7, and can increase calreticulin expression. CD47 blockade simultaneously removes the inhibitory brake (Fig. 1b). Together, these mechanisms enhance macrophage-mediated tumour clearance, as shown in prior NB studies(16).

Here we validate immune competent, chemotherapy-resistant genetically engineered NB models that recapitulate key features of the human NB tumour microenvironment(17). We use these models to test whether ALX301 augments anti-GD2, and whether any added benefit persists when layered onto a clinically aligned temozolomide/irinotecan chemotherapy backbone. We show that CD47 inhibition increases anti-GD2 driven phagocytosis *in vitro* but these effects are not translated *in vivo*, failing to improve anti-GD2 monotherapy or chemoimmunotherapy efficacy in two independent models. Both models exhibit a myeloid-rich, suppressive microenvironment, suggesting that the myeloid niche, not target presence, limits benefit and that macrophage-reprogramming strategies may be required to unlock value from CD47-axis therapy in NB.

## Methods

### *In vivo* establishment of chemoresistant neuroblastoma lines

To model human NB tumours *in vivo*, genetically engineered mouse models (GEMMs) were generated to overexpress MycN and ALK^F1174L^ ligated downstream of the rat tyrosine hydroxylase (*Th*) promoter which developed spontaneous tumours. GEMMs with *Th-MYCN*^+/-^overexpression were used to generate a NB cell line called 9464D, a model which exhibits naturally high resistance to chemotherapy treatment(18). 9464D was further modified to more closely resemble human NB by viral transduction of GD2/GM2 synthase and GD3 synthase to achieve cell surface GD2 presentation and called 9464D-GD2 (gifted by Paul Sondel, University of Wisconsin, US)(19). *Th-MycN* transgenic mice were crossed with *Th-ALK^F1174L^* transgenic mice and the *Th-MycN^+/-^/Th-Alk^F1147L/-^* offspring exhibited higher tumour penetrance and rapid lethality compared to the *Th-MycN* mice(20). Dissociated tumours that were established to become TAM6 and WIN6 were kindly shared by Louis Chesler’s lab at Institute of Cancer Research, Sutton UK. Spontaneous tumours arising in the *Th-MycN^+/-^/Th-Alk^F1147L/-^* 129SvJ/X1 mouse models were subjected to a dose-escalation regimen of VAC chemotherapy (vincristine, adriamycin (doxorubicin), and cyclophosphamide) intraperitoneally to establish a chemotherapy-resistant NB model called TAM6(21). TAM6 tumours were developed by repeated passaging of a spontaneous tumour subcutaneously into another mouse, and each re-established tumour treated with cycles of VAC. The treatment protocol included four cycles with 0.015 mg/kg vincristine, 0.68 mg/kg doxorubicin, and 32 mg/kg cyclophosphamide, followed by two cycles with 0.037 mg/kg vincristine, 1.7 mg/kg doxorubicin, and 80 mg/kg cyclophosphamide. Additionally, a novel chemotherapy-resistant NB model called WIN6 has been developed through a 6^th^ generation backcross of *Th-MycN^+/-^/Th-Alk^F1147L^* 129SvJ/X1 onto C57Bl/6 background. Spontaneous tumours from the *Th-MycN^+/-^/Th-Alk^F1147L^* C57Bl/6 mice were allografted subcutaneously and treated intraperitoneally with a single cycle of VAC chemotherapy calculated at clinically relevant doses: 0.0015mg/kg vincristine, 0.68mg/kg doxorubicin and 32mg/kg cyclophosphamide. Upon tumour regrowth after treatment, the WIN6 model was established as a cell line.

### Cells and culture conditions

The GD2-overexpressing 9464D-GD2 line is maintained under selection with puromycin (6 µg/mL) and blasticidin (7.5 µg/mL)(18,19). Cells grew adherently in DMEM supplemented with 10% FBS, 1% L-glutamine, 1% non-essential amino acids, 1% sodium pyruvate. Neurosphere lines TAM6 (129SvJ/X1) and WIN6 (C57BL/6) were cultured as suspension spheres in DMEM/F-12 supplemented with 15% FCS, B27 (1X), β-mercaptoethanol (0.05 mM), human EGF (0.01 µg/mL) and human bFGF (0.015 µg/mL). Before *in vitro* or *in vivo* use, live cells were enriched by Ficoll separation.

### Animals

All animals were housed in individually ventilated cages with water and food available *ad libitum* and monitored daily. All procedures were reviewed and approved by a University College London (UCL) biological services named animal care and welfare officer and named veterinary surgeon, and performed in accordance with UK Home Office project license PP5675666.

### Therapeutics, concentrations and dosing regimens

Temozolomide (TEM), irinotecan (IRI), and anti-GD2 monoclonal antibody (clone 14G2a, Bio X Cell) were used at clinically relevant levels. Human-equivalent doses (HED) and schedules were adapted from phase II studies in relapsed/refractory NB with patient plasma exposures guiding *in vitro* antibody titrations (0–100 nM)(4). All drugs were administered to mice via intraperitoneal injection throughout. For *in vivo* conversion, human to mouse dose followed Nair *et al.*(22), yielding 100% HED values of 32 mg/kg TEM, 16 mg/kg IRI, and 5 mg/kg anti-GD2. We fixed anti-GD2 at 5 mg/kg and titrated TEM/IRI to identify a suboptimal backbone. We evaluated graded HED backbones at 75% (24/12 mg/kg), 50% (16/8 mg/kg), 25% (8/4 mg/kg), and 10% (3.2/1.6 mg/kg) (TEM/IRI values listed as TEM/IRI). To find the anti-GD2 antibody monotherapy window, we titrated anti-GD2 in a four-dose over two-weeks regimen where high was 2.5 mg/kg, medium was 1.25 mg/kg and low was 0.625 mg/kg per dose. Note the high schedule matches the total anti-GD2 exposure used with TEM/IRI but distributed over two weeks rather than one. The Fc-silent CD47 blocker ALX301 (ALX Oncology, engineered SIRPα D1 fused to mouse IgG1-N297A) was administered at 30 mg/kg, typically four doses over two weeks, per provider guidance. The commercial anti-CD47 antibody (clone MIAP301, Bio X Cell) was used for *in vitro* assays.

### Isolation of bone-marrow derived macrophages (BMDM)

BMDMs were generated *ex vivo* using a method described by Gonzales and Mosser(23). Briefly, femurs from syngeneic mice were flushed with PBS, RBC-lysed using ACK lysis bufffer, and plated on non-tissue-culture-treated 10cm^2^ dishes in DMEM/F-12 (10% FCS, 1% penicillin–streptomycin) with M-CSF (20 ng/mL) for 7 days. At day-7, BMDMs were harvested by treatment with an enzyme-free cell dissociation buffer and were used in downstream assays.

### *In vitro* phagocytosis assays

Tumour cells were labelled with 2uM carboxyfluorescein succinimidyl ester (CFSE) for 20 minutes, washed then mixed with BMDMs at 2:1 (tumour:macrophage) in ultra-low-attachment 96-well plates in serum-free medium. Agents were added as: anti-GD2 antibody alone (0-100nM), CD47 blocker alone (anti-CD47 or ALX301, 0-100nM), or CD47 blocker (0-100nM) plus suboptimal anti-GD2 (0.5nM). After 2 hours at 37 °C, phagocytosis was stopped with two 4°C PBS washes and cells were stained with fixable viability dye (near-IR), CD45 (Pacific Blue) and, for macrophage-only controls, F4/80 (AF700). Phagocytosis was quantified by flow cytometry as percentage CFSE⁺CD45⁺ macrophages and normalised by subtracting the background phagocytosis measured in the control condition in absence of any drug (0nM).

### In vivo studies and treatment combinations

For each treatment arm, n (number of mice) is reported in the figure panels/legends. No formal a priori power calculation was performed because effect sizes in these models were unknown. Group sizes (n = 5-8/arm) were chosen from prior variability and 3Rs principles to detect large, practice-changing effects; key null findings were replicated in independent cohorts and in both models. Mice of 8-12 weeks old were tumour engrafted. To ensure a uniform tumour-bearing cohort at treatment start, an additional ∼40% of mice were engrafted; animals outside prespecified tumour-size limits were not enrolled. In total, 187 mice were engrafted and 115 enrolled and analysed; no treated animals were excluded from analysis.

TAM6 was derived from a tumour in a female mouse and WIN6 from a male mouse; therefore all subsequent allografts were sex-matched within their respective strains to avoid confounding immune responses to sex-linked (e.g., H-Y) antigens.

For subcutaneous engraftment, TAM6, WIN6, or 9464D-GD2 cells were injected into the right flank in 50% Matrigel/50% PBS at optimised doses: 2x10⁵ cells/mouse for TAM6 and 2x10⁶ cells/mouse for 9464D-GD2 (as reported previously(21),(19)); 5x10⁵ cells/mouse for WIN6 yielded consistent growth in C57BL/6J mice. Tumour volume was calculated as (length × width²)/2. Mice were enrolled at ∼10-80 mm³ (unambiguous by calliper measurement). Animals were ranked by baseline volume, binned from smallest to largest, and randomly assigned to groups via a list generator to balance means/SDs across arms.

Temozolomide and irinotecan stocks were stored at −20 °C in DMSO, thawed on ice, diluted in PBS to dose, and co-administered intraperitoneally as a single injection. Anti-GD2 and ALX301 were prepared in PBS, stored at 4 °C at working concentration, and delivered intraperitoneally; when co-dosed, they were mixed and given as a single injection. Each experiment included a matched vehicle-control group: mice received injections on the same schedule as the most intensive regimen, with the appropriate vehicle (PBS or DMSO/PBS) substituted for drug.

### Blood collection and histology

Peripheral blood was collected from the saphenous vein into EDTA tubes and submitted to IDEXX UK for complete blood count (CBC) analysis. For histology during toxicity studies, heart, lung, kidney, liver, spleen, and tumour were harvested after humane euthanasia, fixed in 4% formaldehyde (∼24 h), processed routinely, paraffin-embedded, sectioned, and H&E-stained.

### Humane endpoints and monitoring

Humane endpoints included any single tumour dimension > 15 mm, ulceration at the engraftment site, or treatment-related toxicity defined as > 15% body-weight loss or a moderate grimace score. The primary outcome (had a power calculation been performed) was overall survival, defined as time from treatment start to euthanasia at protocol-defined humane endpoint due to tumour outgrowth. Secondary outcomes included body-weight change and, where performed, CBCs and organ histology. After engraftment, body weight and calliper measurements were recorded twice weekly; animals were observed closely during treatment. Mice were euthanised at humane endpoint or study completion. Survival curves therefore depict protocol-defined overall survival. For all euthanised animals, the reason was tumour burden rather than treatment-related toxicity in the absence of progression. Assessments for euthanasia and calliper measurements were performed by investigators blinded to treatment allocation.

### Tumour microenvironment (TME) flow cytometry

Tumour analysis was done on tumours arising outside treatment enrolment window and were untreated. Untreated tumours were minced with a scalpel, digested with a cell aggregate dissociation medium, Accumax (10 min), and filtered through 70µm strainers. Ammonium chloride potassium (ACK) lysis buffer was used to remove RBCs then cells were split across two panels: an overview panel comprised CD19 (BUV563), NKp46 (BV421), CD11b (BV605), CD45 (FITC), CD3 (PE/Fire700) and GD2 (APC) and a myeloid panel comprised MHCII (BV421), CD11b (BV605), F4/80 (BV711), CD45 (FITC), CD80 (PE), Ly6G (PE/Dazzle594), CD206 (APC) and Ly6C (APC/Cy7). After Fc-block (10 min), viability dye (blue) and antibody master mix were added for 20 min at 4 °C. For intracellular targets, BD Biosciences Perm/Wash^TM^ Buffer was used per instructions before staining (20 min, 4 °C). Samples were fixed and run on a BD Symphony within 3 days. Analysis was performed in FlowJo. The gating strategy is shown in Supplementary Fig. S1.

### Statistical analysis

Data was processed in Microsoft Excel and graphed in GraphPad Prism. Each point represents a biological replicate. Bars show mean ± s.d. unless otherwise indicated. Group comparisons used two-way ANOVA with Tukey’s multiple comparisons correction; exact P values are shown for significant contrasts. Survival analyses (where applicable) used log-rank (Mantel–Cox) tests.

## Results

### Establishment and immune contexture of chemoresistant, immune competent NB models

NB is an embryonal tumour derived from neural crest progenitor cells. *MYCN* gene amplification and *ALK* activating mutations are associated in promoting aggressive tumours in ultra-high-risk NB patients(24,25). To model these tumours *in vivo*, genetically engineered mouse models (GEMMs) have been generated to overexpress MycN and ALK^F1174L^ downstream of the rat tyrosine hydroxylase (*Th*) promoter(20,26). Genes downstream of the *Th* promoter are expressed in committed sympathetic precursor cells of the neural crest. Expression of oncogenes *Th*-*MycN* and *Th-ALK^F1174L^* give rise to tumours in the sympathetic ganglia and adrenal gland, locations commonly seen in human NB. We generated chemoresistant NB cell lines from *Th-MYCN/Th-ALK^F1174L^* spontaneous tumours by sequential addition of vincristine, doxorubicin and cyclophosphamide to establish TAM6 cells on 129SvJ background (previously described(21)), and new model WIN6 on C57BL/6 background for syngeneic use. A *Th-MYCN* derived 9464D line on C57BL/6 background was transduced to overexpress GD2 (9464D-GD2) for selected comparisons(19) (Fig. 2a).

**Figure 2.**
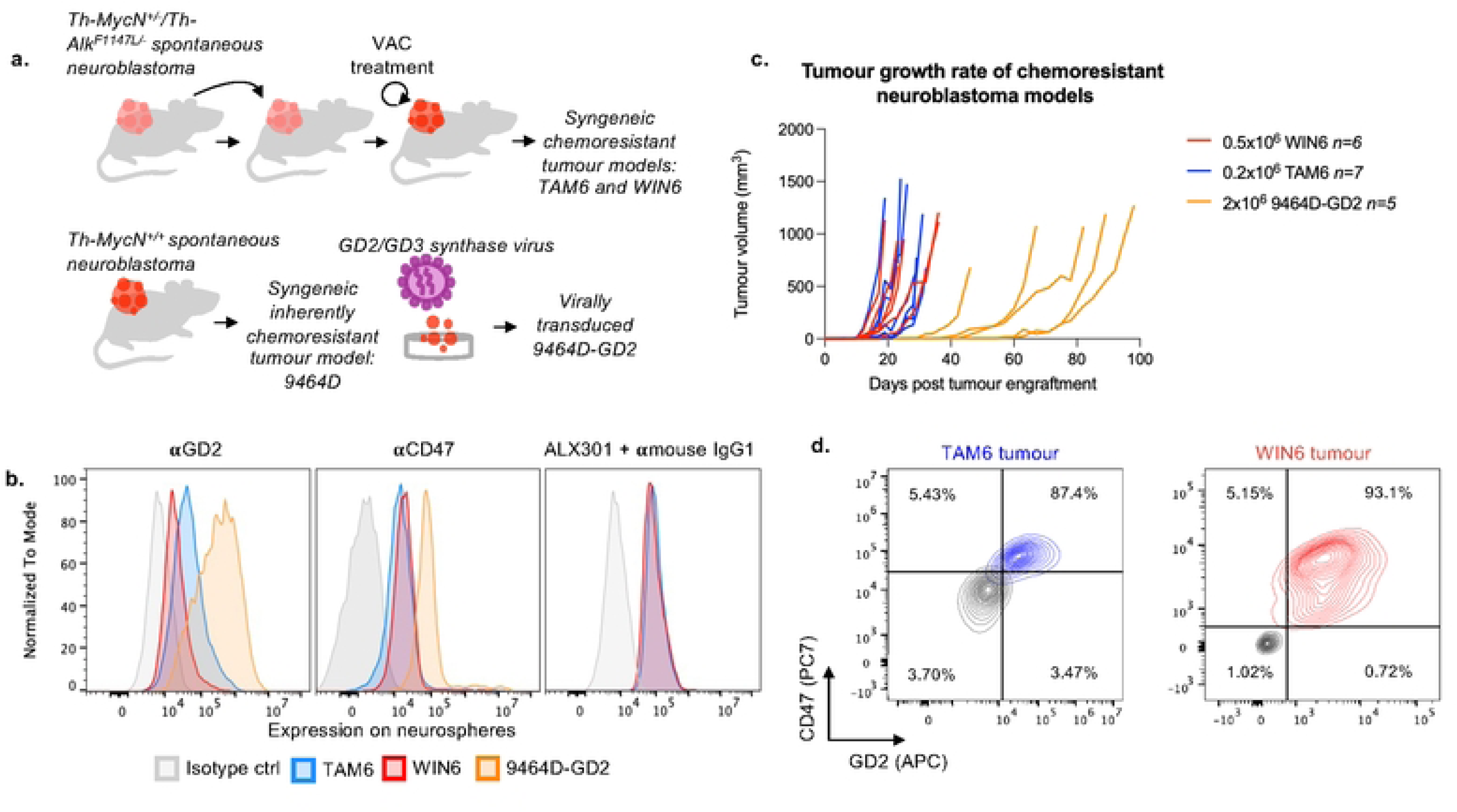
Establishment and characterization of immunocompetent, chemoresistant neuroblastoma models. **a.** Schematic of chemoresistant neuroblastoma cell line generation; TAM6, WIN6 and 9464D-GD2. **b.** Flow cytometry confirms target antigen expression on TAM6, WIN6 and 9464D-GD2 cells, showing surface GD2 and CD47. ALX301 binding to tumour cells was detected using secondary anti-mouse IgG1 staining. **c.** Tumour growth kinetics of subcutaneously engrafted TAM6, WIN6 and 9464D-GD2 cells. **d.** Flow cytometric contour plots to demonstrate freshly dissected and stained TAM6 and WIN6 tumours also express GD2 and CD47.

By flow cytometry, TAM6 and WIN6 expressed GD2 and CD47, and ALX301 bound tumour cells (Fig. 2b). Optimised subcutaneous engraftment yielded reproducible growth for all models although 9464D-GD2 grew slower than TAM6 or WIN6 (Fig. 2c). Dissected and freshly stained TAM6 and WIN6 tumours confirmed *in vivo* GD2 and CD47 expression (Fig. 2d).

High-parameter flow cytometry showed that TAM6 tumours harbour a significantly larger myeloid compartment, whereas WIN6 tumours are relatively lymphoid-rich; both models contain immunosuppressive myeloid populations, including MDSCs and M2-like macrophages (Fig. 3a, b; gating in Supplementary Fig. S1).

**Figure 3.**
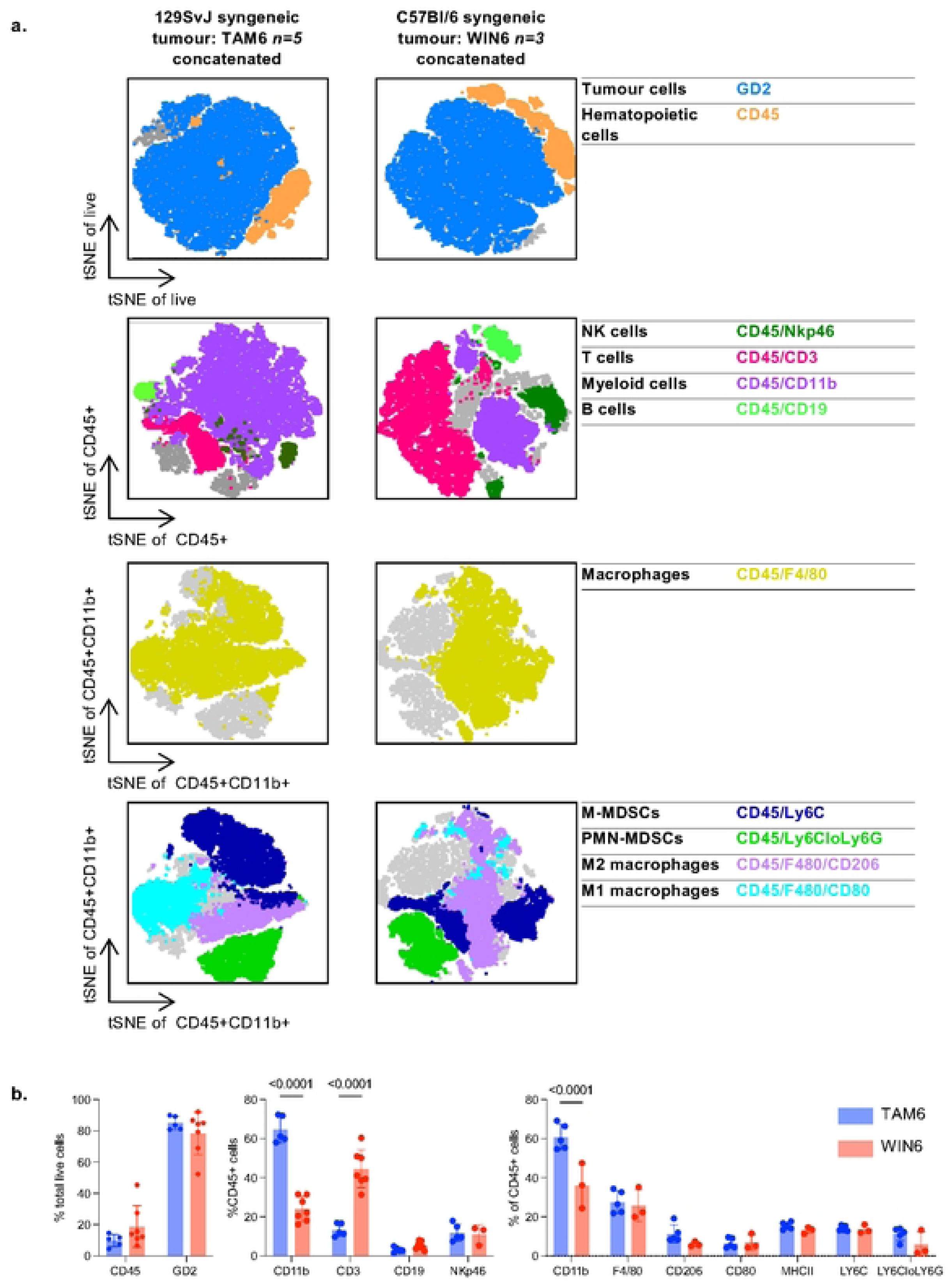
Immune profiling of TAM6 and WIN6 tumours. **a.** Immune profiling of TAM6 and WIN6 tumours by high-parameter flow cytometry and dimensionality reduction by t-Distributed Stochastic Neighbour Embedding (t-SNE) analysis. **b.** TAM6 tumours displayed a significantly larger myeloid compartment within the CD45 gated cells, compared with WIN6.

### CD47 blockade augments anti-GD2 mediated phagocytosis *in vitro*

To test whether CD47 blockade augments anti-GD2 antibody induced ADCP in these chemoresistant models, CFSE-labelled tumour cells were co-cultured *in vitro* with syngeneic bone marrow derived macrophages at 2:1 tumour:macrophage ratio for 2 hours. Anti-GD2 alone induced dose-dependent phagocytosis in both TAM6 and WIN6 models, whereas ALX301 or anti-CD47 (MIAP301) alone did not increase uptake above baseline. Combining either form of CD47 blockade with suboptimal anti-GD2 (0.5nM) produced a clear additive increase in phagocytosis of TAM6 and WIN6 across biological replicates (Fig. 4a, b). Based on the percentage uptake of CFSE, the TAM6 model is approximately 5 fold less susceptible to phagocytosis than the WIN6 model (Fig. 4b). By contrast, 9464D-GD2, despite high GD2 and CD47 cell surface expression, was resistant to anti-GD2 ± CD47 blockade in the same assay, even when 10-fold greater concentration of anti-GD2 was used, despite its being susceptible to ADCC in prior work(19) (Fig. 4c, d). As 9464D-GD2 was resistant to ADCP, but not ADCC, we tested whether conditioned-media from 9464D-GD2 cultures inhibited anti-GD2 ADCP of WIN6 cells. It did not, indicating that soluble inhibitors released by these 9464D-GD2 cells were not the mechanism behind the resistance of this tumour to ADCP in these *in vitro* assays (Fig. 4e).

**Figure 4.**
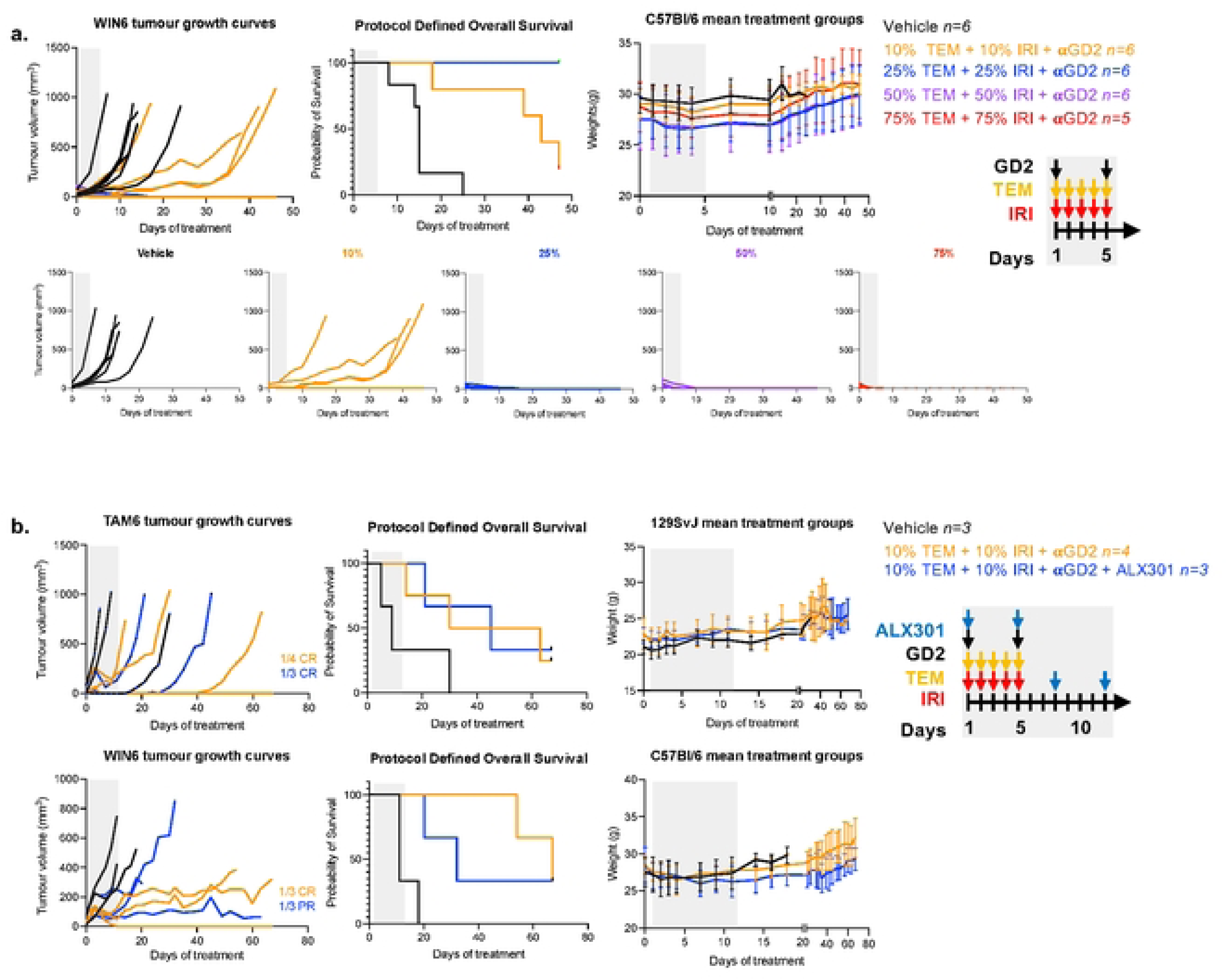
CD47 blockade enhances anti-GD2 mediated phagocytosis in vitro of TAM6 and WIN6 tumour lines. **a.** Representative flow cytometry plots of phagocytosis assays. Two populations can be seen, tumour cells (CFSE population) and syngeneic bone marrow–derived macrophages (CD45 population). Macrophage uptake of tumour cells was quantified as CFSE+CD45+ events indicated by the orange box**. b.** Quantification of phagocytosis across independent biological replicates (n=3 per model). Suboptimal anti-GD2 antibody (0.5nM) was combined with CD47 blockade using ALX301 or the commercial anti-CD47 antibody. **c.** In vitro phagocytosis assays with 9464D-GD2 tumour cells and syngeneic macrophages. **d.** Increasing anti-GD2 concentration tenfold did not overcome resistance in 9464D-GD2 cells. **e.** Phagocytosis of WIN6 tumour cells was unaffected by conditioned supernatant from 9464D-GD2. Error bars represent mean ± SD; statistical significance between three sources of macrophages was determined using two-way ANOVA with Tukey’s multiple comparison test.

### CD47 blockade does not improve anti-GD2 antibody or chemoimmunotherapy *in vivo*

As CD47 blockade augmented ADCP *in vitro* by anti-GD2 for TAM6 and WIN6 tumour cells, we next asked whether this would translate to anti-tumour effects *in vivo.* We first modelled the clinically used chemoimmunotherapy schedule by dosing temozolomide and irinotecan (TEM/IRI) on consecutive days 1-5 with anti-GD2 on days 1 and 5. Dose titration of TEM/IRI identified a partially efficacious regimen at 10% the human-equivalent dose (HED), TEM/IRI (3.2 mg/kg / 1.6 mg/kg) with anti-GD2 (5 mg/kg) in the WIN6 model, which produced delayed growth without cures (Fig. 5a). Due to its proposed superior toxicity profile compared with anti-CD47 antibodies, we selected ALX301 as a CD47 targeting agent. Adding ALX301 (30 mg/kg) to this chemoimmunotherapy backbone did not improve tumour control or survival in either WIN6 or TAM6 models (Fig. 5b).

**Figure 5.**
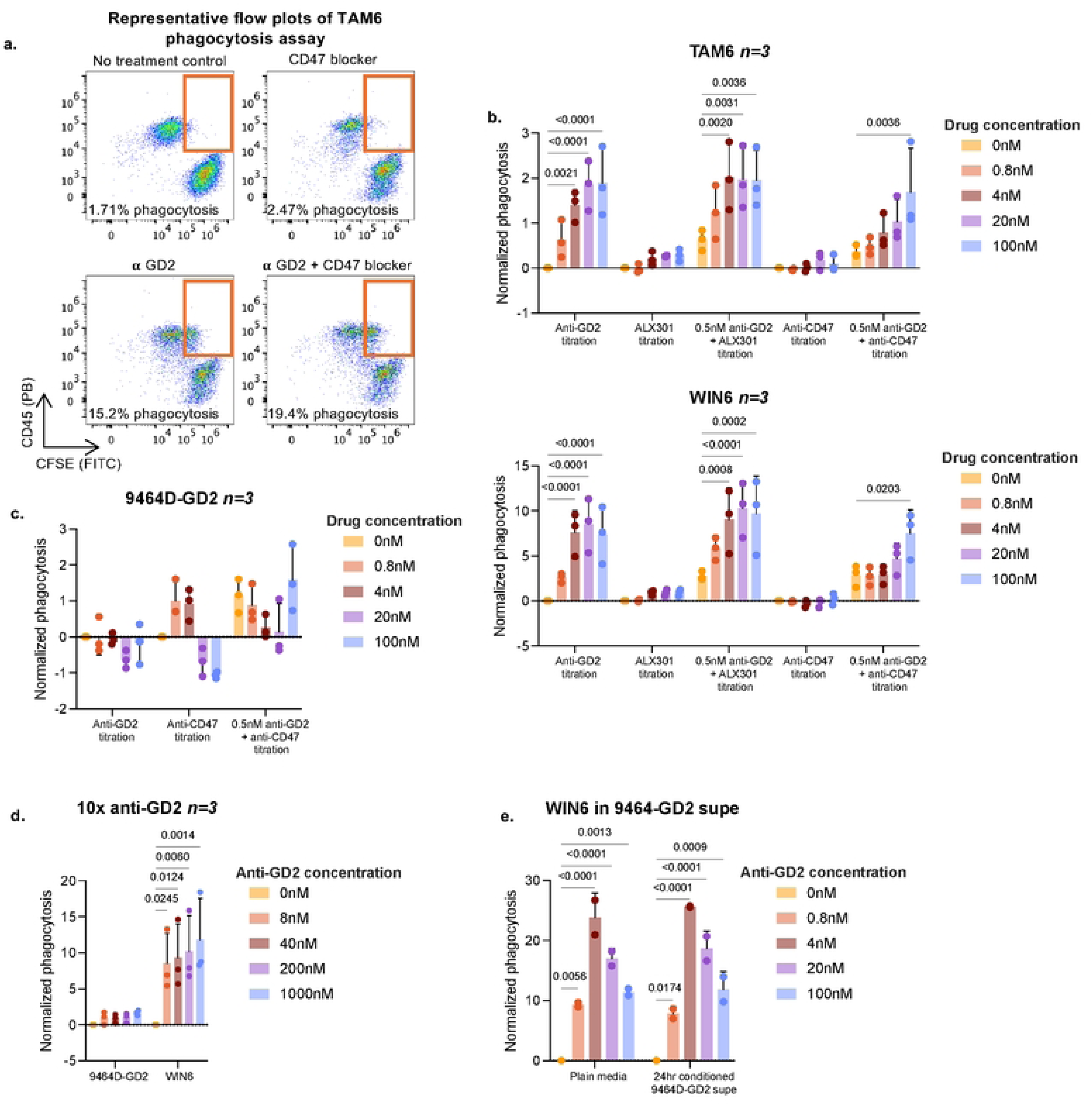
Initial analyses of ALX301 in combination with partially effective chemoimmunotherapy. **a.** WIN6 tumour growth curves, survival, and mean treatment group weights following dose titration of temozolomide (TEM) plus irinotecan (IRI) in combination with anti-GD2 (5 mg/kg). Percentages shown are that of human equivalent dose (HED) for TEM and IRI. **b.** Tumour growth curves, survival, and mean treatment group weights are shown for combination of ALX301 (30mg/kg) with the partially efficacious chemoimmunotherapy regimen in both WIN6 and TAM6 models. Treatment schemas are shown on the right.

Recognizing that ALX301 was not designed for monotherapy use, we hypothesized that a regimen with prolonged anti-GD2 treatment to allow ALX301 and anti-GD2 co-administration might be necessary to achieve the synergistic effect required for macrophage activation and effective tumour cell clearance. An anti-GD2 monotherapy (1.25 mg/kg) dosed four times over two weeks was partially efficacious in both WIN6 and TAM6 models (Suppl. Fig. S2). However, when co-administrated with ALX301 (30 mg/kg), no additional efficacy was observed compared to the effects of anti-GD2 alone, again suggesting a lack of synergy between the two therapies in WIN6 and TAM6 models under these conditions (Fig. 6a, b).

**Figure 6.**
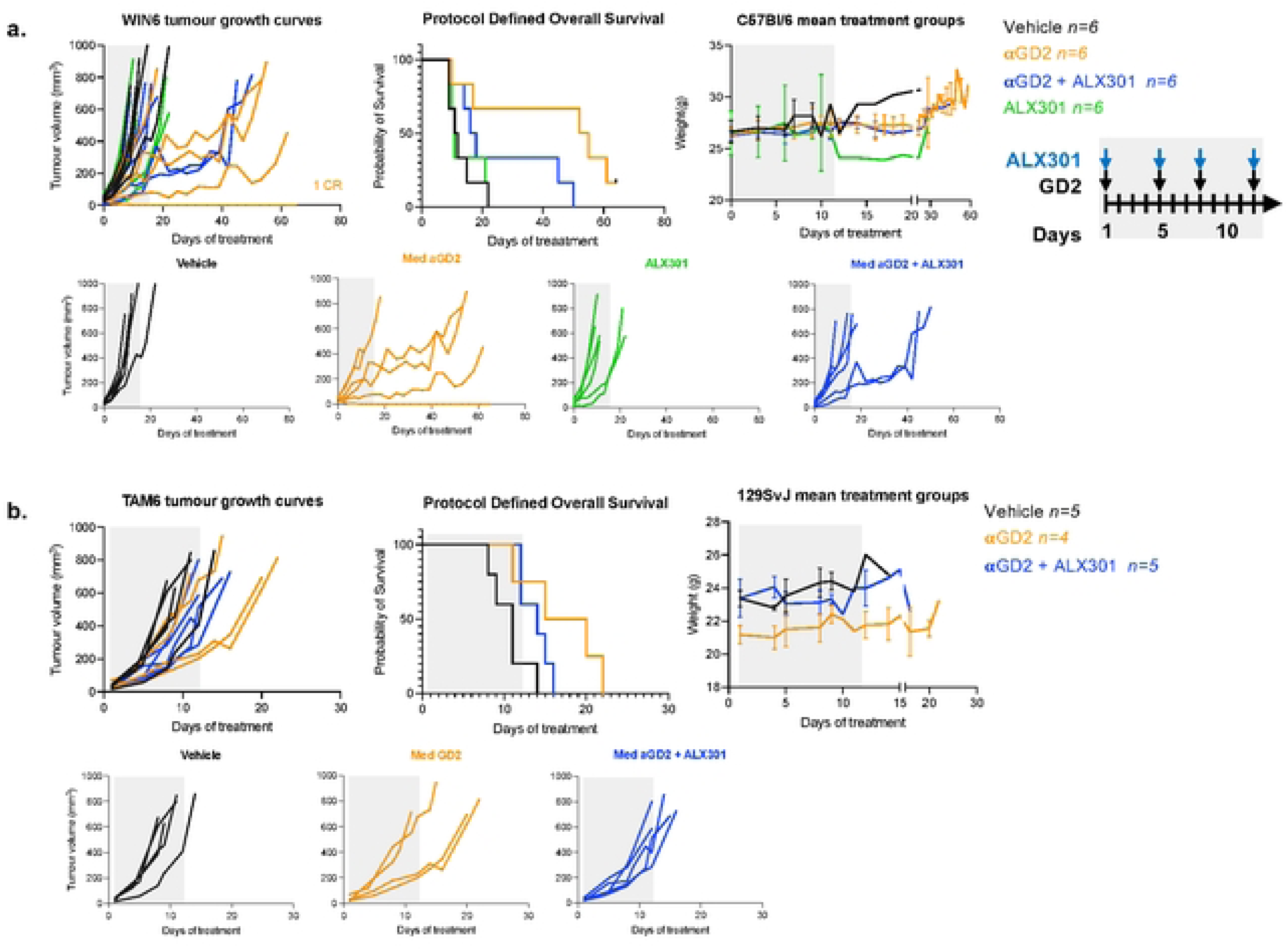
CD47 blockade does not enhance the efficacy of suboptimal anti-GD2 monotherapy in vivo. **a.** WIN6 tumour growth curves, survival, and mean treatment group weights following treatment with vehicle, suboptimal anti-GD2 (1.25 mg/kg), ALX301 (30 mg/kg), or the combination of anti-GD2 plus ALX301. Anti-GD2 suboptimal dose finding results are shown in supplementary figure 2. **b.** Equivalent experiment in TAM6 tumours, with treatment groups as indicated. Treatment schedule is shown on the right.

### Intensified CD47 blockade with chemoimmunotherapy showed favourable safety profile but no additional benefit

To investigate whether increased exposure could unmask synergy, we extended treatment with an additional two doses of anti-GD2 and ALX301 layered onto the TEM/IRI backbone in the WIN6 model. Efficacy remained unchanged with ALX301 addition (Fig. 7a). Complete blood counts (CBCs) were obtained at the end of treatment on day 19 or at the humane endpoint for animals with rapidly growing tumours requiring euthanasia prior to the day 19 end of treatment-scheduled blood sampling. These revealed small but significant decreases in haemoglobin and haematocrit (RBC %) with CD47 blockade, consistent with on-target RBC effects, without effects on clinical condition, grimace scores, or body weight (weights summarised as mean ± SD in Fig. 7a) (Fig. 7b). Histopathology (H&E) of heart, lung, kidney, liver, spleen, and tumour showed no tissue injury (Fig. 7c). These findings suggest that while this regimen of ALX301 does not appear to introduce systemic toxicity when combined with the chemoimmunotherapy backbone, within these two immune competent neuroblastoma models, it also does not enhance the efficacy of the treatment compared to the chemoimmunotherapy backbone alone.

**Figure 7.**
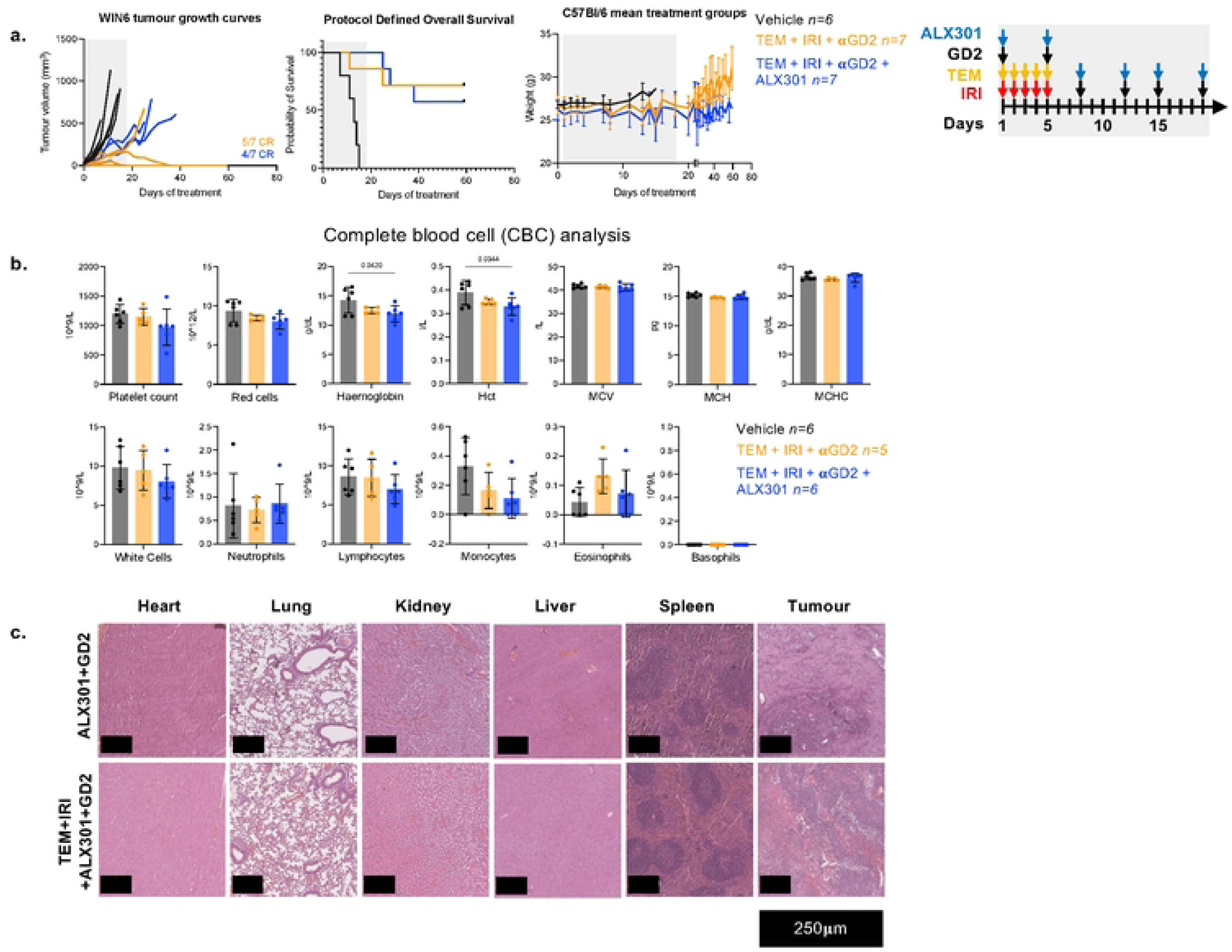
Impact of chemotherapy addition to anti-GD2 and ALX301 on toxicity and efficacy. **a.** WIN6 tumour growth curves, survival, and mean treatment group weights following an extended anti-GD2 and ALX301 treatment schedule combined with chemotherapy (as in Fig. 5). **b.** To assess toxicity, peripheral blood samples were collected from treated mice either prior to tumour-related death or at the end of treatment, and complete blood counts (CBCs) were performed. **c.** Histological analysis of haematoxylin and eosin (H&E) stained sections from heart, lung, kidney, liver, spleen, and tumours. Representative images are shown at 5× magnification, with scale bar representing 250 µm.

## Discussion

We provide a translational stress test of CD47 blockade in combination with anti-GD2 immunotherapy or chemoimmunotherapy in immune competent, chemoresistant NB mouse models. A key contribution of this study is the introduction and immune mapping of two immunocompetent, chemoresistant neuroblastoma models, WIN6 (C57BL/6) and TAM6 (129SvJ). Both models express the clinical target GD2 and the inhibitory ligand CD47, engraft reproducibly, and recapitulate the myeloid-rich, immunosuppressive niche seen in human neuroblastoma, including abundant MDSCs and M2-like macrophages (with TAM6 particularly myeloid-dominant). These features make WIN6 and TAM6 useful beyond the CD47 question: they enable clinically aligned dosing studies and testing of new combinations, and their intact TME allows interrogation of cell-cell interactions by flow, spatial, and single-cell readouts. The observation that CD47 blockade did not enhance anti-GD2 antibody or chemoimmunotherapy in these stringent models is informative. Together, WIN6 and TAM6 provide a rigorous, disease-relevant platform to derisk combinations, define mechanisms of resistance, and support translational applicability.

Using macrophages derived from bone marrow of healthy mice, an Fc-silent CD47 inhibitor (ALX301) enhanced anti-GD2 driven phagocytosis *in vitro*, confirming target biology at the macrophage-tumour synapse in two independent models, WIN6 and TAM6. However, in the same syngeneic immune competent models, no *in vivo* benefit emerged when ALX301 was added to anti-GD2 or to a clinically aligned TEM/IRI with anti-GD2 regimen. Safety was acceptable and small, with expected RBC-related changes without clinical or histologic toxicity, implicating insufficient efficacy rather than tolerability as the barrier in these models. The doses and route of administration of ALX301 in TAM6 and WIN6 are those previously shown to effect tumour responses in other models(16). The augmentation of phagocytosis *in vitro* used the same batch of reagent as the *in vivo* studies excluding the possibility of batch failure. Mechanistically, our models are myeloid-heavy and suppressive, with abundant MDSCs and M2-like macrophages, particularly in TAM6. In such niches, removing the CD47 “Don’t Eat Me” signal alone may be necessary but not sufficient to trigger productive macrophage-mediated tumour clearance. The 9464D-GD2 findings of resistance to phagocytosis further highlight cell-intrinsic resistance; our *in vitro* analysis indicates this resistance is not mediated by soluble factors and is not rescued by higher anti-GD2 dose, underscoring context-dependence of the macrophage response.

These findings contrast with the strong synergy reported by Theruvath and co-workers using the same anti-GD2 (clone 14G2a) and ALX301(16). In that study, synergy was shown in one syngeneic model derived from a different GEMM (TH-MYCN in 129/SvJ mice) and in several human xenografts in immunodeficient hosts. Our results therefore indicate model-dependent heterogeneity: the regimen that was sufficient in those systems was not in WIN6 or TAM6. It remains possible that higher ALX301 exposure could provide benefit here; we did not escalate beyond the prolonged schedule in Fig. 7 because the dose/regimen used has documented activity and reflects provider guidance.

These observations are consistent with broader clinical experience for CD47-axis agents: manageable on-target hematologic effects but limited efficacy without a conducive microenvironment(27,28). Our data suggest the field would benefit from investigating rational myeloid reprogramming to unlock CD47 blockade capabilities. There are several therapies already validated for preconditioning myeloid cells in the context of antitumour therapies, included are CSF1R blockade (to reduce suppressive macrophages)(29,30), CD40 agonists or STING/TLR agonists (to license APCs and skew macrophages)(31–33) or low-dose radiation (to boost “Eat Me” cues)(34). On-treatment TME profiling to ascertain myeloid states and antigen presentation may be crucial to pinpoint where CD47 might contribute.

Strengths of this study include two immunocompetent, chemoresistant models with detailed immune contexture, clinically aligned dosing/schedules, matched *in vitro* and *in vivo* testing, and integrated safety readouts. Next steps should include on-treatment TME sampling and tests of myeloid-conditioning combinations with optimised scheduling; if tolerated, increased ALX301 exposure could also be explored. These efforts will clarify if, and under what conditions, CD47 blockade adds value in neuroblastoma.

## Conclusion

In our neuroblastoma models, this ALX301 CD47 blockade agent, used in this regimen, does not augment anti-GD2 or chemoimmunotherapy *in vivo*, despite clear *in vitro* activity and acceptable safety. We propose that subsequent preclinical studies should test whether adding a myeloid-reprogramming agent to this regimen may create a pro-inflammatory macrophage niche to enable CD47 blockade to augment the efficacy of anti-GD2 chemoimmunotherapy.

## Acknowledgements

We would like to acknowledge grant support from Solving Kids Cancer, National institute of Health Research, Little Princess Trust, Cancer Research UK Grand Challenge, SU2C, Great Ormond Street Hosp Biomedical Research Centre and Research into Chidhood Cancer (RICC).

